# Critical capillary waves of biomolecular condensates

**DOI:** 10.1101/2023.10.29.564316

**Authors:** Shunsuke F. Shimobayashi, Paul J. Ackerman, Tomo Kurimura, Takashi Taniguchi, Clifford P. Brangwynne

## Abstract

Biomolecular condensates formed by phase separation are key players in cellular organization, yet their interfacial mechanics remain poorly understood. Here, we show that both synthetic and endogenous nuclear condensates exhibit critical-like interfacial behaviors near the phase boundary, including enhanced capillary fluctuations, critical slowing down, and reduced surface tension. By combining optogenetic control with sub-micron-resolution fluctuation spectroscopy, we quantitatively estimate surface tension, bending rigidity, and effective viscosity. Surface tension diminishes as the system approaches the critical composition, consistent with classical theories of phase separation. Notably, bending elasticity emerges as an unexpected feature of these nuclear liquid-like structures, suggesting the formation of structured interfacial layers that progressively weaken near criticality. Among these condensates, the nucleolus displayed exceptionally high viscosity, which may arise in part from viscoelastic coupling to the surrounding perinucleolar heterochromatin, effectively increasing the apparent viscosity in the long-time fluctuation regime. This non-invasive approach enables probing condensate mechanics in living cells and may provide a basis for diagnosing or modulating condensates in biomedical contexts.

## I. INTRODUCTION

Biomolecular condensates formed via liquid-liquid phase separation (LLPS) have emerged as a fundamental mechanism for organizing the intracellular environment without membranes [1–6]. These dynamic compartments concentrate specific proteins and nucleic acids to facilitate diverse biological processes, including gene expression, RNA metabolism, and signal transduction. Importantly, aberrant condensate formation has been linked to a range of diseases, such as neurodegenerative disorders and cancers, highlighting the need to understand the physical principles governing their material properties [7–9]. Among these properties, surface tension plays a central role in defining condensate shape, stability, and dynamics, particularly during processes such as fusion or wetting [10, 11]. Bending rigidity—typically considered irrelevant for liquid-like droplets—has recently been suggested to contribute to condensate morphology [12], raising the possibility that condensates possess unexpectedly complex interfacial mechanics. A central open question is how condensate interfacial mechanics behave throughout the phase diagram, particularly near a critical point. Given that intracellular condensates are inherently multicomponent, non-equilibrium systems, it remains unclear whether their interfacial behavior near criticality conforms to theoretical expectations established for classical critical fluids [13].

## II. RESULTS

### A. Quantification of surface fluctuations and material properties of biomolecular condensates in living cells

To quantify the interface properties of condensates in living cells, we first examine a tractable optogenetic system, Corelets [3, 14]. Corelets enable control of phase separation through light-activated oligomerization of proteins including the intrinsically disordered regions (IDRs) of FUS or HNRNPA1 (FUS_N_, 1–214; HNRNPA1_C_, 186–320), model aromatic-rich polar sequences in human osteosarcoma (U2OS) cells [3, 14] (Fig. 1a). We activate optogenetic oligomerization and then record time-lapse images of the resulting condensates and detect the contours (see methods), clearly observing surface fluctuations with a velocity of approximately 10 nm per sec (Fig. 1b,c). We find that the time-averaged contours are generally not perfect circles (Fig. 1b), likely due to the inhomogeneous mechanical environment in living cells. In order to quantify the fluctuations in the condensate equatorial plane, we consider the contour deviation *δr*(*φ,* Δ*t*) with a time-lag Δ*t*

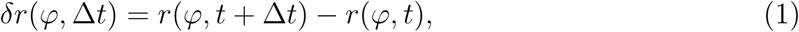

where we set a polar coordinate system from the center of mass of each condensate (Fig. 1b). Fig. 1d shows the fluctuation amplitude ⟨|*δr*(Δ*t*)|^2^⟩ of condensate surfaces as a function of lag time Δ*t*. For both FUS_N_- and HNRNPA1_C_-Corelets, the fluctuation amplitude increases with Δ*t* and saturates to a plateau. To quantify these dynamics, the fluctuation amplitude |*δr*(*φ,* Δ*t*)|^2^ was first averaged over all time point pairs with the same Δ*t* within a single condensate, yielding ⟨|*δr*(*φ,* Δ*t*)|^2^⟩, and then subsequently averaged over the angular coordinate *φ* to obtain ⟨|*δr*(Δ*t*)|^2^⟩. The decay toward the plateau can be well described by a single exponential, ⟨|*δr*(Δ*t*)|^2^⟩ ∝ 1 − *e*^−Δ^*^t/^*^Δ^*^t^*^0^, from which we determined relaxation times Δ*t*_0_ of 2.6 ± 1.3 s and 1.1 ± 0.1 s for FUS_N_- and HNRNPA1_C_-Corelets, respectively. Fourier transformation of experimentally measured fluctuations ⟨|*δr*(*φ,* Δ*t*)|^2^⟩ yields the ensemble average of the fluctuation spectrum, that is, ⟨|Δ*B_m_*(Δ*t*)|^2^⟩ with mode number *m*, which will appear later in Eq. (4).

**FIG. 1.**
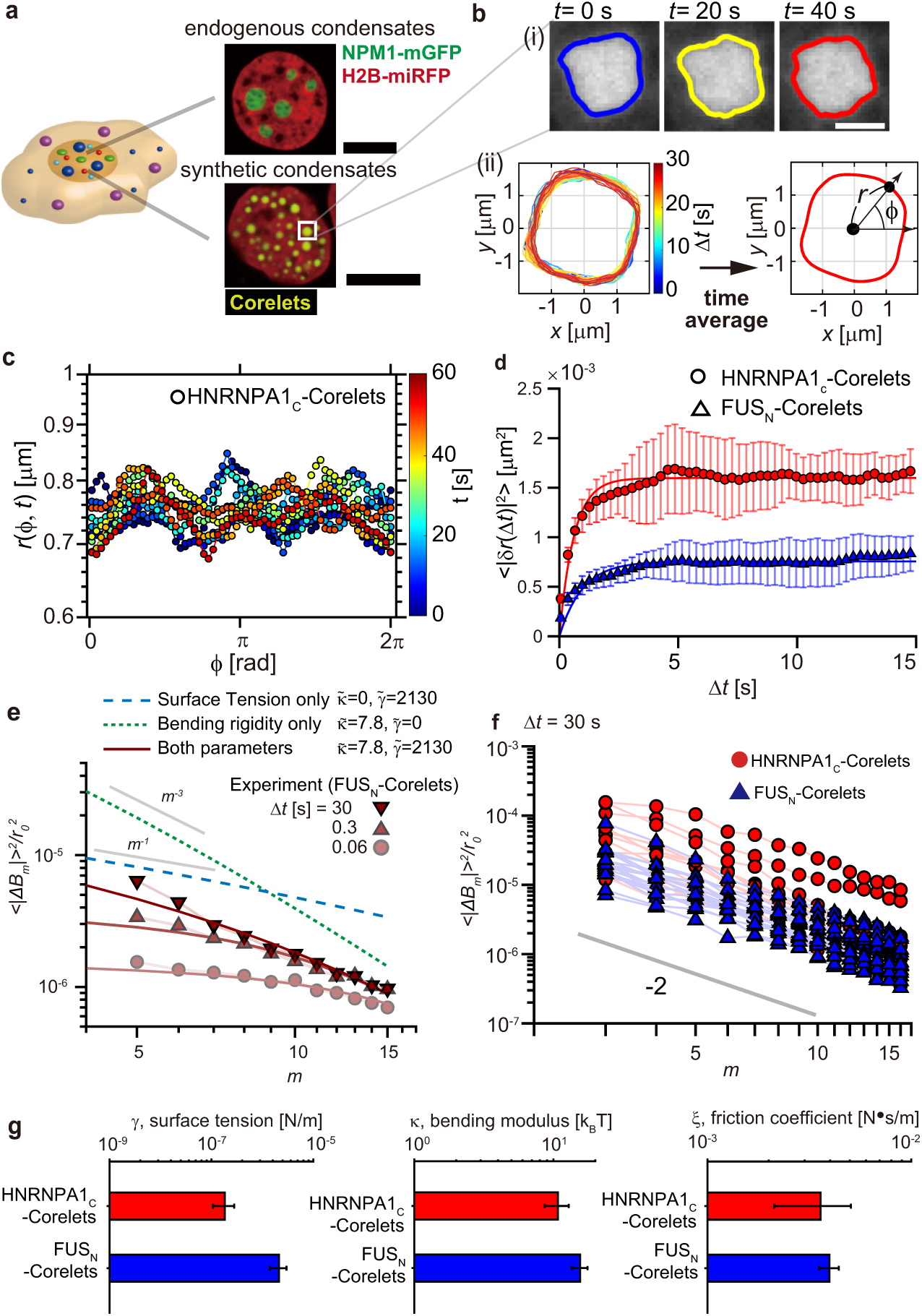
Quantification of surface fluctuations and material properties of biomolecular condensates in living cells. (a) Confocal fluorescence images of U2OS cells show nucleoli (NPM1-mGFP) and synthetic condensates (HNRNPA1_C_-Corelets). Scale bar, 10 *µ*m. (b)(i) Surface fluctuation of HNRNPA1_C_-Corelets. The condensate interfaces are represented by the solid lines. Scale bar, 1 *µ*m. (ii) Overplots and time-average of contours of HNRNPA1_C_-Corelets over 30 sec with Δ*t* = 180 ms (167 frames) (c) Temporal dynamics of condensate contour *r*(*φ, t*) for HNRNPA1_C_-Corelets (*t* = 0, 12, 24, 36, 48, 60 s). (d) Ensemble average of fluctuation amplitude ⟨|*δr*(Δ*t*)|^2^⟩ as a function of Δ*t* for FUS_N_ and HNRNPA1_C_-Corelets. The solid lines show the best fit to ⟨|*δr*(Δ*t*)|^2^⟩ ∝ 1 −*e*^−Δ*t/*Δ*t*0^. (e) Normalized fluctuation spectrum of FUS_N_-Corelets against the mode number *m*. The experimental data (symbols) are shown for different time lags (Δ*t* = 30, 0.3, and 0.06 s). The solid lines represent the best fits to the data using Eq. (4). The dashed lines exhibit a power law with a slope of *m*^−3^ for the surface tension-only model and *m*^−1^ for the bending rigidity-only model. (f) Normalized fluctuation spectrum against *m* with Δ*t* = 30 s for FUS_N_ (*n* = 21) and HNRNPA1_C_-Corelets (*n* = 14). (g) Box plots comparing surface tension (left) and bending modulus (right), with red and blue boxes representing HNRNPA1-Corelets and FUS_N_-Corelets, respectively. The error bars represent the standard error, respectively. *n* is the number of analyzed condensates.

To theoretically elucidate the fluctuation behavior, we model the free energy functional of the condensate surfaces, which can be expressed as

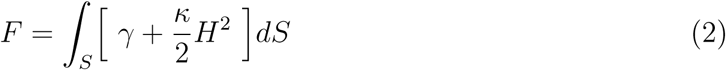

here *γ*, *κ* and *H* are condensate surface tension, surface bending modulus, and surface local curvature. The equation of motion of the condensate surface can be expressed as;

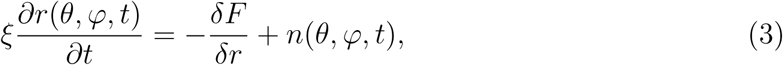

where *ξ* is a friction coefficient and *n*(*θ, ϕ, t*) is a noise term reflecting microscopic fluctuations (e.g. thermal noise). Assuming thermally-driven surface fluctuations on the cross-sectional plane at *θ* = *π/*2 in the spherical coordinate system (See Supplemental Figure S1), the fluctuation spectrum ⟨|Δ*B_m_*(Δ*t*)|^2^⟩ can be expressed as;

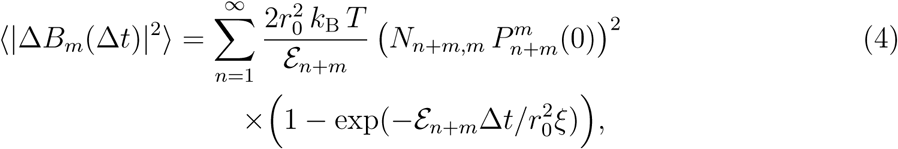

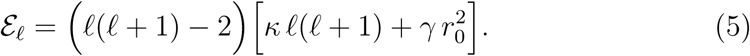

Here, 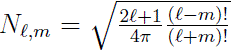 and 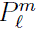 is the associated Legendre polynomial. The parameters *r*_0_, *k*_B_, *T*, *ξ* are the mean condensate radius in the equatorial plane, the Boltzmann constant, temperature, and friction coefficient, respectively. The parameters *γ*, *κ* and *ξ* can be measured by fitting ⟨|Δ*B_m_*(Δ*t*)|^2^⟩ with Eq. (4). For our fit, modes (*m* ≥ 15) were excluded because the analysis approaches the resolution limit of the method and the spectra are increasingly affected by measurement noise, though still above the noise floor [15].

Fig. 1e shows the representative fluctuation spectrum for a single condensate of FUS_N_-Corelets over 500 frames at time lags Δ*t* = 0.06, 0.3, and 30 s with the best-fit lines obtained using Eq. (4). As shown in the supplemental figure S2, the spectrum for Δ*t* = 30 s can be considered as the Δ*t* → ∞ case for FUS_N_-Corelets. In this limit, the spectrum can be fitted using the expression 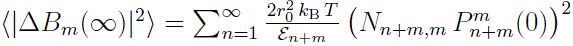, yielding *γ* and *κ* (Fig. 1e). The spectrum for Δ*t <* 30 s was fitted by Eq. (4), where the values of *γ* and *κ* obtained from the previous fitting were substituted, to determine *ξ*. The theoretical prediction, as shown by dashed lines, demonstrates that the *κ* contribution exhibits an *m*^−3^ scaling, while the *γ* contribution exhibits an *m*^−1^ scaling. Hence, the −2 power law, as experimentally observed, signifies a regime where both terms contribute concurrently. Moreover, Fig. 1f shows the fluctuation spectrum ⟨|*B_m_*(Δ*t*)|^2^⟩ for a large set of individual FUS_N_ and HNRNPA1_C_ condensates (500 frames, Δ*t* = 30 s), exhibiting a power law with a slope of nearly −2 against *m*. Fitting yields statistical values for a bending rigidity *κ* ranging from 1 to 10 k_B_*T*, consistent with previous estimates for stress granules [12]. Surface tension *γ* ranges from 10^−7^ to 10^−5^ N/m, in alignment with previously reported values for both in vitro and in vivo condensates [11, 12, 16–18]. The extracted friction coefficient *ξ* spans 10^−3^ to 10^−2^ N·s/m; to our knowledge, this represents the first quantitative measurement of interfacial friction for intracellular condensates.

### B. Quantification of surface fluctuations and material properties of endogenous nuclear condensates

Next, we aimed to test the applicability of our approach to endogenous nuclear condensates, focusing on nucleoli—sites of ribosomal biogenesis, and nuclear speckles, which serve as hubs for mRNA processing and splicing factors [11, 19] (Fig. 2a). As shown in Fig. 2b, the two condensates exhibited distinct magnitudes and time scales of interfacial fluctuations. We computed the fluctuation spectra ⟨|Δ*B_m_*(Δ*t*)|^2^⟩ with Δ*t* = 750 s for nucleoli and Δ*t* = 250 s for nuclear speckles, revealing a power-law decay with an exponent of −2 and confirming the presence of larger fluctuations in nuclear speckles (Fig. 2c). To rule out experimental noise, we performed identical measurements after fixation with 4% paraformaldehyde, verifying that the observed fluctuation signals were well above the noise floor, particularly for long-wavelength modes (low *m*) (Supplemental Fig. S3).

**FIG. 2.**
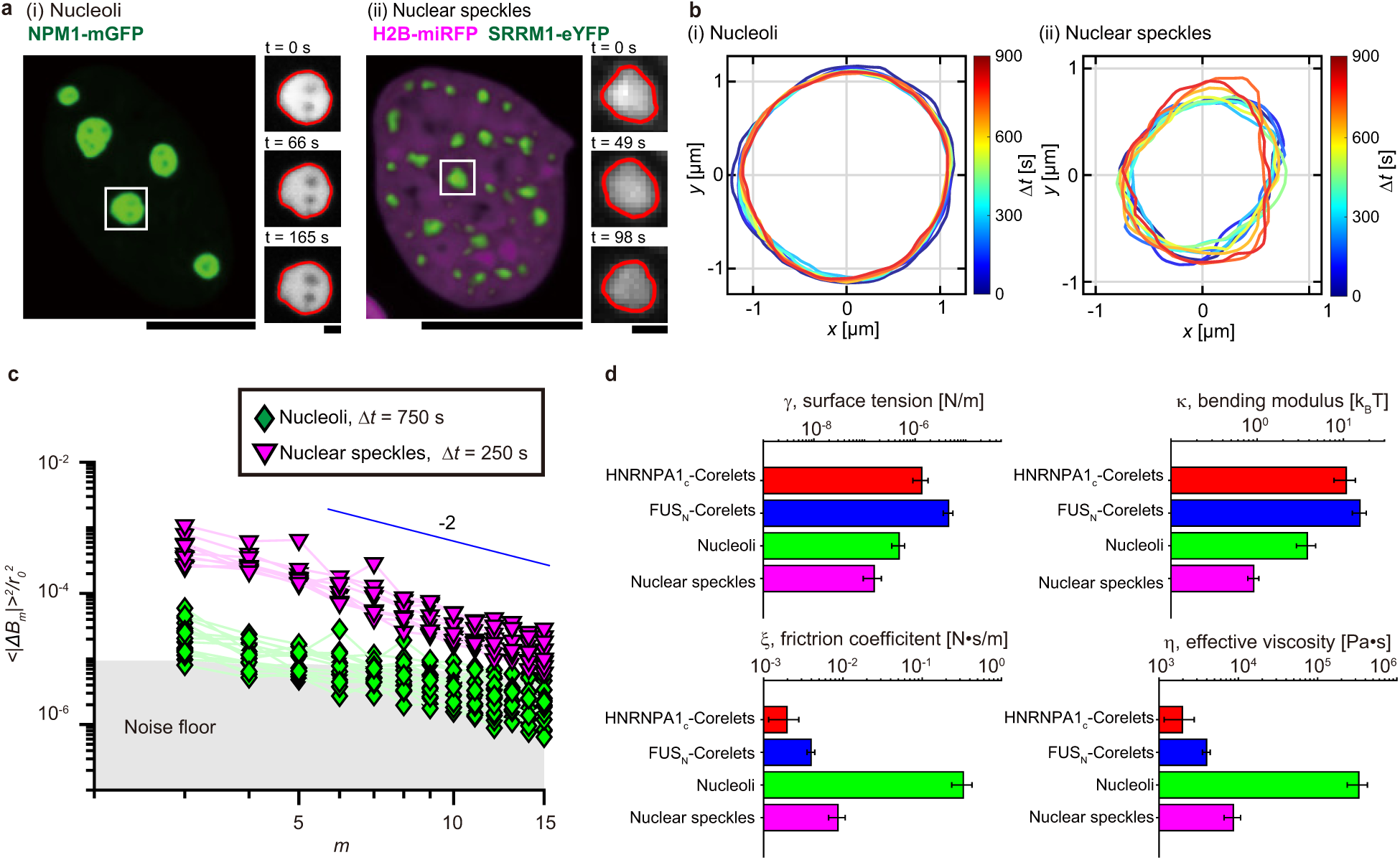
Quantification of surface fluctuations and material properties of endogenous nuclear condensates. (a) Fluorescence images of U2OS cells show nucleoli (NPM1-mGFP) and nuclear speckles (SRRM1-eYFP). Time-lapse confocal images of a single condensate are also displayed. (b) Overplots of a single condensate contour of (i) nucleoli and (ii) nuclear speckles in U2OS cells with (i) Δ*t* = 30 s (50 frames) and (ii) Δ*t* = 10 s (50 frames). (c) Normalized fluctuation spectrum against the mode number *m*with Δ*t* = 750 sec for nucleoli (*n* = 15) and Δ*t* = 250 sec for nuclear speckles (*n* = 19). (d) Bar graphs show the mean and standard error of surface tension (*γ*), bending modulus (*κ*), friction coefficients (*ξ*) and effective viscosity (*η*) for HNRNPA1-Corelets (*n* = 14), FUS_N_-Corelets (*n* = 21), nucleoli (*n* = 15), and nuclear speckles (*n* = 9), respectively. Scale bar, 10 *µ*m. *n* is the number of analyzed condensates.

From these spectra, we determined the surface tension, bending modulus, friction coefficient, and effective viscosity as summarized in Fig. 2d. The surface tension of nuclear condensates ranged from 10^−7^ to 10^−6^ N/m, consistent with prior measurements in HeLa cells and *Xenopus* oocytes [11, 16], as well as colloid-polymer emulsions [20]. Strikingly, the bending modulus of these endogenous condensates again fell within 1–10 *k*_B_*T*, suggesting that intracellular condensates—both nuclear and cytoplasmic, native and synthetic—may share similar levels of interfacial stiffness. Given the multicomponent nature of intracellular condensates, it is plausible that certain components preferentially accumulate at the interface, forming a compositionally distinct and structured layer. Such a layer may exhibit lateral molecular organization and cohesive interactions, akin to a Langmuir monolayer [10], thereby contributing to the observed bending rigidity of the interface. Supporting this notion, recent work on *C. elegans* P granules has demonstrated that MEG-3 forms low-dynamic protein clusters at the condensate interface to suppress coarsening and enhance structural integrity [21]. These findings raise the possibility that membrane-less organelles may exhibit membrane-like mechanical properties due to surface-bound molecular scaffolds. Since our focus is on nuclear condensates, it is also tempting to speculate that such scaffolds could be provided by the surrounding chromatin matrix. However, their contribution would likely manifest in the apparent viscosity rather than in the bending rigidity, as our analysis probes long-time fluctuations beyond the relaxation regime. Notably, the nucleolus, despite being surrounded by perinucleolar heterochromatin, did not show a substantially higher *κ* than other nuclear condensates (Fig. 2d).

In contrast, nucleoli exhibited an anomalously high friction coefficient, over an order of magnitude greater than that of Corelet condensates and nuclear speckles (Fig. 2d). Given that the low-mode regime is dominated by surface tension rather than elastic restoring forces, and that viscous dissipation is expected to dominate over elastic losses, the friction coefficient *ξ* can be related to the effective viscosity *η* via *ξ* ≈ *ηr* [22], where *r* denotes the condensate radius. Using the experimentally measured *r*, we can evaluate the effective viscosity, shown in a bar graph (Fig. 2d). The effective viscosities we obtained for Corelet condensates and nuclear speckles are comparable to previously reported large-scale viscosities of the surrounding nucleoplasm (10^2^–10^3^ Pa·s) measured by microrheology and magnetic bead experiments [23, 24]. This is also approximately a thousand times higher than those of most *in vitro* condensates (∼1 Pa·s), highlighting the highly viscous intracellular environment [17, 18].

Notably, the viscosity of nucleoli reaches ∼ 10^5^ Pa·s, which is about a hundred times greater than that of the nucleoplasm. Such exceptionally high viscosity highlights the distinct physical state of the nucleolus among nuclear condensates, characterized by a highly dense, multiphase organization with RNA- and RNP-rich subcompartments [11, 25, 26], which is also reflected by their distinct refractive index [27]. In addition, the contribution of perinucleolar heterochromatin likely elevates the apparent viscosity: any viscoelastic coupling from the surrounding chromatin *η* ≈ *Gτ*, where *G* is the shear modulus and *τ* is the structural relaxation time, would be extracted as a viscous rather than an elastic component in the long-time regime [28]. For comparison, previous studies that measured nucleoli from *Xenopus* oocytes reported viscosities on the order of 10^2^–10^3^ Pa·s based on fusion dynamics [11] and micropipette aspiration [29]. Those are about two orders of magnitude lower than the effective viscosity we estimated for nucleoli, possibly because these amphibian germinal vesicle nucleoli lack the surrounding perinucleolar heterochromatin.

### C. Analysis of critical capillary waves in synthetic versus endogenous condensates

To gain further physical insights into the material properties of condensates, we leveraged controllable optogenetics to explore their critical behavior. By adjusting blue-light intensity, we could precisely modulate the binding affinity between core and IDR components, thereby tuning the system’s position within the phase diagram. As the light intensity was decreased, the system approached the critical point, i.e., *χ* → *χ_c_* and *C*_dil_*/C*_den_ → 1 (Fig. 3a). The Flory-Huggins theory predicts that surface tension vanishes at the critical point [30]. Consistent with this, we observed a marked increase in surface fluctuations (Fig. 3b,c) and a slowdown in fusion dynamics (Fig. 3d) near the critical point, reflecting enhanced capillary fluctuations and critical slowing down [20]. By fitting the fluctuation spectra ⟨|Δ*B_m_*(Δ*t*)|^2^⟩, we quantified *γ* and *κ* across a range of concentration ratios. Consistent with theoretical expectations, surface tension decreased as *C*_dil_*/C*_den_ → 1 (Fig. 3g). In contrast, the bending rigidity also decreased, but this behavior likely reflects structural changes at the interface, such as the loss of interfacial organization or adsorption, rather than a universal critical behavior (Fig. 3e).

**FIG. 3.**
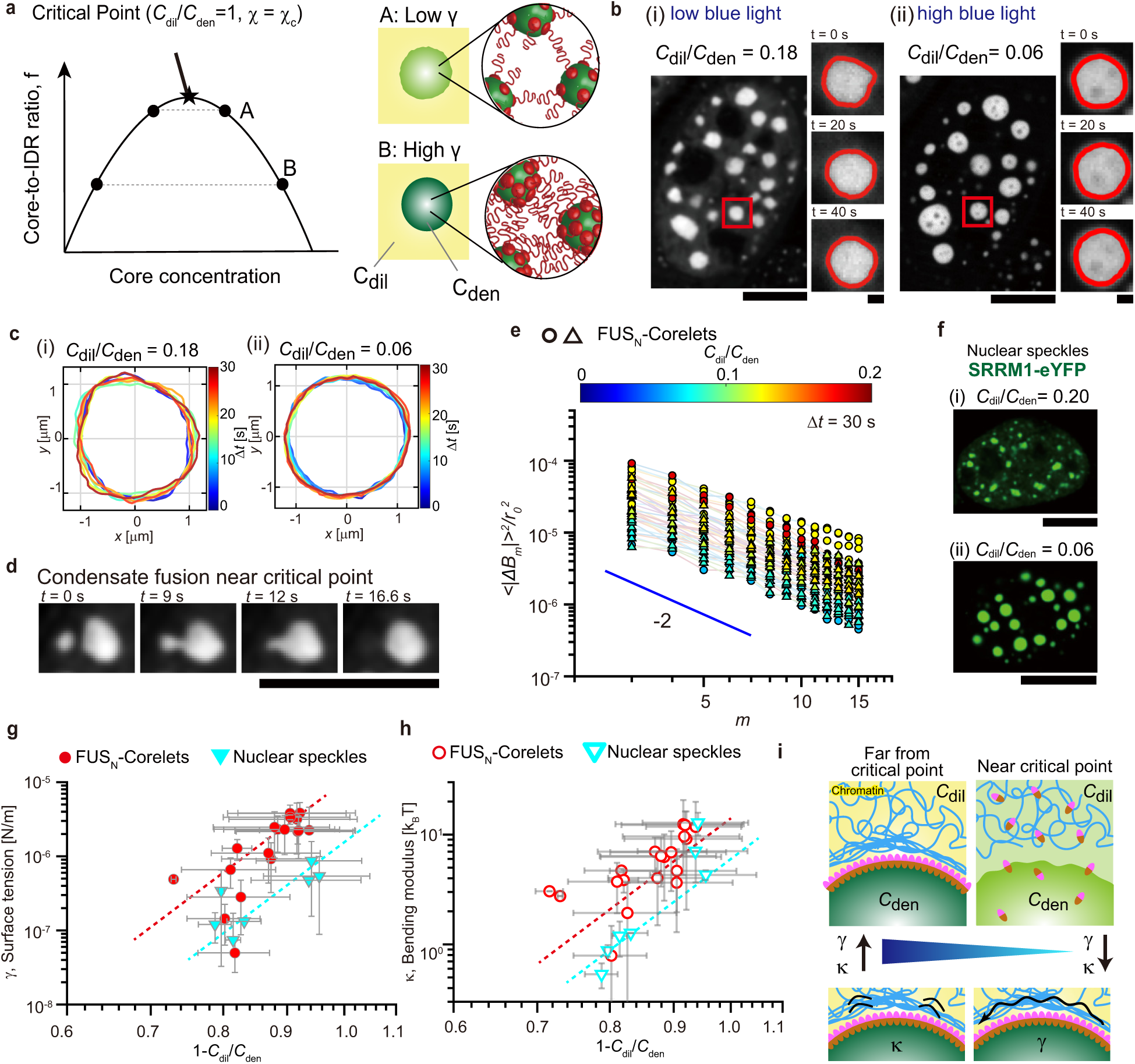
Analysis of critical capillary waves in synthetic versus endogenous condensates. (a) Schematic of a phase diagram of Corelet system shows that surface tension *γ* decreases as a cell approaches the critical point, where *χ* = *χ_c_* and *C*_dil_*/C*_den_ = 1. (b) Fluorescence images of U2OS cells expressing FUS_N_-Corelets subjected to approximately (i) 10^−4^ W/cm^2^ and (ii) 10^−2^ W/cm^2^ intensities of blue light. Scale bar, 10 *µ*m. Time-lapse confocal images of a single condensate are also displayed. Scale bar, 1 *µ*m. (c) Overplots of single condensate contours in U2OS cells expressing FUS_N_-corelets under approximately (i) 10^−4^ W/cm^2^ and (ii) 10^−2^ W/cm^2^ intensities of blue light with Δ*t* = 180 ms over 30 sec. (d) A droplet fusion of FUS_N_-corelet condenates in a U2OS cell near the critical point. (e) Fluctuation spectrum ⟨|Δ*B_m_*(Δ*t*)|^2^⟩*/r*^2^ against *m* with Δ*t* = 30 s and various ratios *C*_dil_*/C*_den_ for U2OS cells expressing FUS_N_-Corelets (*n* = 60). (f) Fluorescence images of U2OS cells expressing different levels of SRRM1-eYFP. (g, h) Surface tension *γ* and bending modulus *κ* plotted against 1 − *C*_dil_*/C*_den_ for FUS_N_-Corelets (103 condensates from 19 cells) and nuclear speckles (49 condensates from 13 cells). Data points represent single-cell means; error bars indicate standard deviations. The dashed lines are eye-guides. (i) Schematic illustration of condensate interfaces far from (left) and near (right) the critical point.

To assess whether similar critical behavior exists in endogenous condensates, we overexpressed SRRM1—a scaffold protein of nuclear speckles—beyond its native levels. This led to more spherical speckles, lower *C*_dil_*/C*_den_, and reduced interfacial fluctuations (Fig. 3f,g), accompanied by a clear increase in surface tension. Despite the complex composition of endogenous condensates, surface tension exhibited similar critical behavior to synthetic systems (Fig. 3g), reinforcing the view that critical phenomena underlie both classes.

## III. DISCUSSION AND PERSPECTIVE

Our findings reveal that nuclear condensates exhibit critical-like interfacial behavior, including the vanishing of surface tension near the phase boundary—consistent with classical theories of phase transitions. Beyond surface tension, our fluctuation spectroscopy uncovered bending rigidity as an unexpectedly prominent feature of these nominally liquid-like structures, suggesting the presence of structured interfacial layers stabilized by preferential adsorption or lateral organization of specific molecular components. Bending rigidity progressively decreased as the system approached criticality (Fig. 3h,i), likely reflecting the loss of such interfacial structure. Moreover, we observed striking variations in effective viscosity: most nuclear condensates exhibited values of 10^2^–10^3^ Pa·s, whereas the *nucleolus* reached ∼ 10^5^ Pa·s, possibly reflecting viscoelastic coupling to the surrounding *perinucleolar heterochromatin*.

Taken together, our study integrates optogenetic control with non-invasive fluctuation spectroscopy to quantify the material properties of condensates—including surface tension, bending rigidity, and viscosity—across synthetic and endogenous contexts. The ability to resolve these parameters at the sub-micron scale opens new opportunities to relate condensate material properties to biological function and dysfunction, enabling biophysical probing of pathological condensates through their intrinsic fluctuations and informing rational strategies to target aberrant phase behavior.

## ACKNOWLEDGMENTS

We wish to acknowledge Kota Konishi, Akira Miyaki, Nguyen Thai Le and Masumi Sanada for experimental and analytical support. This work was supported in parts by JST, PRESTO Grant Number JPMJPR21E8 (to S.F.S.), Japan, JSPS KAKENHI Grant Numbers 18K13521, 23K13072, 25H01328 (to S.F.S.), research grants from iPS Cell Research Fund (to S.F.S.), the Uehara Memorial Foundation (to S.F.S.), Nakatani foundation (to S.F.S.), The Kao Foundation for Arts and Sciences (to S.F.S.), 2022 iPS Academia Japan Grant (to S.F.S.), JST FOREST Program, Grant Number JPMJFR230V (to S.F.S.), AFOSR MURI grant (to C.P.B.), and Princeton MRSEC (to C.P.B.).

## AUTHOR CONTRIBUTIONS

S.F.S., T.T. and C.P.B. designed the research; S.F.S. and P.J.A. developed the analysis codes; S.F.S. performed the experiments; S.F.S. and T. K. analyzed the experimental and numerical data; S.F.S. wrote the manuscript; S.F.S, P.J.A., T.T., T. K., and C.P.B. reviewed and edited the draft.

