## Supplemental materials for "Critical capillary waves of biomolecular condensates"

Shunsuke F. Shimobayashi*
*Center for iPS Cell Research and Application, Kyoto University, Sakyo-ku, Kyoto 606-8507, Japan*

Paul J. Ackerman
*Department of Chemical and Biological Engineering,*
*Princeton University, Princeton, NJ 08544, USA*

Tomo Kurimura
*Center for iPS Cell Research and Application, Kyoto University, Sakyo-ku, Kyoto 606-8507, Japan*

Takashi Taniguchi
*Department of Chemical Engineering, Kyoto University, Kyoto 615-8510, Japan*

Clifford P. Brangwynne
*Department of Chemical and Biological Engineering,*
*Princeton University, Princeton, NJ 08544, USA*
*Howard Hughes Medical Institute, Princeton University, Princeton, NJ 08544, USA and*
*Omenn-Darling Bioengineering Institute, Princeton University, Princeton, NJ 08544, USA.*
(Dated: December 4, 2025)

#### CONTENTS

I. Methods 2
A. Cell culture 2
B. Plasmid construction 2
C. Lentiviral transduction 2
D. Construction of stable cell lines 2
E. Corelets (Core scaffolds to promote droplets) 2
F. Cell fixing 2
G. Microscopy 3
H. Free energy of condensate surface 3
I. Fluctuation dynamics of condensate surface 4
J. Fluctuation analysis of condensate surface 5
K. Quantification of concentration ratio 6

II. Figures 7

References 8

*

### I. METHODS

#### A. Cell culture

Human cell lines used in this study include U2OS and Lenti-X 293T (Takara Bio USA). Lenti-X 293T cells were only used for virus production, while U2OS cells were used for experiments. Cells were cultured in growth medium (DMEM, Nacalai) supplemented with 10% FBS (Wako) and penicillin and streptomycin (Nacalai) at 37°C with 5% CO2 in a humidified incubator.

#### B. Plasmid construction

Plasmids were transformed into Stellar cells (Clontech), from which single colonies were picked, grown in LB supplemented by Ampicillin for 16 hours, and miniprepped (QIAGEN) following manufacturer instruction. All cloning products were confirmed by sequencing (GENEWIZ). The constructs (pHR-FUS$_{\mathrm{N}}$-mCherry-sspb, pHR-HNRNPA1$_{\mathrm{C}}$-mCherry-sspb, pHR-NLS-iLID-EGFP-FTH1, FM5-NPM1-GFP, pHR-H2B-miRFP670, pHR-mGFP-P2A-mCherry, FM5-SRRM1-eYFP) are derived from those used in our previous studies [1, 2].

#### C. Lentiviral transduction

Lentiviruses were produced by cotransfecting Lenti-X 293T cells grown to approximately 70% confluency, controlled with Countess 3 FL (Thermo), in 6-well plates with transfer plasmids (1.5 µg), pCMV-dR8.91 (1.33 µg) and pMD2.G (0.17 µg) using FuGENE HD Transfection Reagent (Promega) following manufacturer's instruction. After 2 days, 2 mL of supernatant containing viral particles was harvested and filtered with 0.45 µm filter (Merck). Supernatant was immediately used for transduction or stored at -80°C in aliquots.

#### D. Construction of stable cell lines

U2OS cells were grown to 10-20% confluency, controlled with Countess 3 FL (Thermo), on 96 well glass-bottom dishes (Cellvis) and 10-100 µl of filtered viral supernatant was added to the cells. Virus-containing medium was replaced with fresh growth medium 48 hours post-infection. The cells were typically imaged no earlier than 72 hr after infection.

#### E. Corelets (Core scaffolds to promote droplets)

Corelets have a two-module optogenetic system that mimics the endogenous oligomerization of IDR-rich proteins to drive biomolecular condensation, using a light-activatable high valency core [1, 2]. The core is comprised of 24 human ferritin heavy chain (FTH1) protein subunits, which self-assemble to form a spherical particle of 12 nm diameter (referred to as "Core"), which is fused to a nuclear localization signal (NLS) and an engineered protein iLID. The second module is comprised of a self-interacting IDR fused to SspB, which upon blue light activation strongly heterodimerize ($K_{\mathrm{d}} \sim 130$ nM) with iLID to form self-interacting particles.

#### F. Cell fixing

U2OS Cells stably expressing NPM1-mGFP were fixed with 4% paraformaldehyde (Nacalai) in PBS (Nacalai) and incubated for 12 min at room temperature, followed by three times rinse with PBS.

##### G. Microscopy

Fluorescence images were taken using a spinning-disk confocal microscope (Yokokgawa CSU-W1) with Nikon 100x oil immersion objective (CFI Plan Apo λD, NA 1.45) and an Andor iXon Ultra 888 EMCCD camera on a Nikon Eclipse Ti2-E body. Samples were maintained at 37°C and 5% CO2 with a stage top incubator (Tokai Hit). 488, 561, and 640 nm lasers (Stradus) were used for imaging mGFP/eYFP, mCherry, and miRFP, respectively. 488 nm laser light was also used for activating iLID at an excitation power of approximately $10^{-4}$ to $10^{-2}$ W/cm$^2$ as measured with a microscope slide photodiode power sensor (PM100D, Thorlabs). Phase separation dynamics for the Corelet system were captured by imaging in mCherry channel with global activation by 488 nm laser. All image acquisition was performed using Nikon NIS-Elements AR software. All image analysis was performed using custom-built Python and MATLAB scripts (The Math Works).

##### H. Free energy of condensate surface

We consider the surface fluctuations of incompressible condensates with a volume $V$. The free energy of the condensate is given by the sum of the surface energy and the bending energy as

$$F = \int_S \Big[ \frac{\kappa}{2} H^2 + \gamma \Big] dS \tag{1}$$

where $\kappa$ is the bending rigidity, $H/2$ the mean curvature of the surface, $\gamma$ the surface tension[3, 4]. The integrals are performed over the condensate surface. Because of the incompressibility of the system, the total volume of the condensate $V$ is constant, $V = V_0$. Using a polar coordinate system, a point on the condensate surface at a time $t$ is expressed by $\boldsymbol{r}(\theta,\varphi,t) = r(\theta,\varphi,t)\boldsymbol{e}_r$, where $r(\theta,\varphi,t)$ is the radius of the condensate and $\boldsymbol{e}_r$ is the radial unit vector. $r(\theta,\varphi,t)$ is expressed as

$$r(\theta,\varphi,t) = r_0 + R(\theta,\varphi,t), \tag{2}$$

$$R(\theta,\varphi,t) = C_0 + \sum_{\ell=1}^{\infty} \sum_{m=-\ell}^{\ell} C_{\ell m}(t) Y_{\ell m}(\theta,\varphi) \tag{3}$$

where $r_0 \equiv (3V_0/4\pi)^{1/3}$, $Y_{\ell m}$ is the spherical harmonics, $C_{\ell m}(t)$ is the complex expansion coefficients which have a relation $C^*_{\ell m} = C_{\ell,-m}$ so that $R$ is real. Now we consider a small shape fluctuation of a condensate from a spherical shape with a radius $r_0$, that is, $|R| \ll r_0$. Substituting equations (2) and (3) into eq.(1), and expanding the free energy in terms of $|C_{\ell m}|$ up to the second order, the free energy is expressed as

$$\Delta F = F - F_0 = \sum_{\ell=2}^{\infty} \sum_{m=-\ell}^{\ell} \frac{\mathcal{E}_\ell}{2r_0^2} |C_{\ell m}|^2 \tag{4}$$

$$\mathcal{E}_\ell \equiv \big(\ell(\ell+1) - 2\big) \Big[\kappa\ell(\ell+1) + \gamma r_0^2\Big] \tag{5}$$

where $F_0 \equiv 8\pi\kappa$, and $C_0 = -(r_0/4\pi)\sum_{\ell m} |C_{\ell m}|^2$ is used to satisfy the volume conservation up to the 2nd order in $C_{\ell m}$. The probability that a shape fluctuation with $\{C_{\ell m}\}$ occurs can be given by the Boltzmann factor as

$$P(\{C_{\ell m}\}) \propto \exp\Big[ - \frac{\Delta F(\{C_{\ell m}\})}{k_{\mathrm{B}} T} \Big], \tag{6}$$

where $k_{\mathrm{B}}$ and $T$ are the Boltzmann constant and temperature, respectively. From eq.(6),

$$\langle |C_{\ell m}|^2 \rangle = \frac{2 r_0^2 k_{\mathrm{B}} T}{\mathcal{E}_\ell} \tag{7}$$

#### I. Fluctuation dynamics of condensate surface

The equation of motion of a condensate surface can expressed as;

$$\xi \frac{\partial r(\theta,\varphi,t)}{\partial t} = -\frac{\delta F}{\delta r} + n(\theta,\varphi,t), \tag{8}$$

where $\xi$ is a friction coefficient and $n(\theta,\phi,t)$ is a external noise. Here, we consider a thermal noise that satisfies the following fluctuation-dissipation theorem:

$$\langle n(\mathbf{\Omega},t)n(\mathbf{\Omega}',t')\rangle = \frac{2k_{\mathrm{B}}T}{\xi}\delta(t-t')\delta(\mathbf{\Omega}-\mathbf{\Omega}')/\sqrt{g} \tag{9}$$

where $\mathbf{\Omega}=(\theta,\varphi)$ and $g$ is the metric of the surface. From eq.(8), the equation for the mode $C_{\ell m}(t)$ is written as

$$\frac{dC_{\ell m}(t)}{dt} = -\Gamma_\ell C_{\ell m}(t) + n_{\ell m}(t), \tag{10}$$

where $\Gamma_\ell$ is defined as $\Gamma_\ell = \mathcal{E}_\ell/(r_0^2\xi)$. $n_{\ell m}$ is defined as $n_{\ell m} \equiv \int n(\Omega,t)Y_{\ell m}(\Omega)\sqrt{g}d\theta d\varphi$ and from eq.(9) it satisfies the following relation:

$$\langle n_{\ell m}(\mathbf{\Omega},t)n_{\ell' m'}(\mathbf{\Omega}',t')\rangle = \frac{2k_{\mathrm{B}}T}{\xi}\delta(t-t')\delta_{\ell\ell'}\delta_{mm'} \tag{11}$$

In the experiment, the two-dimensional shape of the condensate view from $z=\infty$ is observed as

$$r(\theta=\pi/2,\varphi,t) = r_0 + \sum_{m=-\infty}^{\infty} B_m(t)e^{im\varphi} \tag{12}$$

It should be noted that $r(\varphi,t)$ in eq.(12) means $r(\theta=\pi/2,\varphi,t)$. Then, the correlation of $\delta r(\varphi,t) \equiv r(\varphi,t)-r(\varphi,0)$ between two points on the contour, $\langle \delta r(\varphi,t)\delta r(\varphi',t)\rangle$, is evaluated.

$$\langle \delta r(\varphi,t)\delta r(\varphi',t)\rangle = \sum_{m=-\infty}^{\infty} \langle|\Delta B_m(t)|^2\rangle e^{im(\varphi-\varphi')} \tag{13}$$

where $\Delta B_m(t) \equiv B_m(t) - B_m(0)$.

When using eqs.(2)-(3), the correlation can be expressed as that of $\delta r(\varphi,t) \equiv R(\theta=\pi/2,\varphi,t) - R(\theta=\pi/2,\varphi,0)$. Namely,

$$\begin{aligned}\langle \delta r(\varphi,t)\delta r(\varphi',t)\rangle = &\sum_{\ell=1}^{\infty}\sum_{m=-\ell}^{\ell} 2\big(1-e^{-\Gamma_\ell t}\big)\\ &\times Y_{\ell m}(\frac{\pi}{2},\varphi)Y_{\ell,-m}(\frac{\pi}{2},\varphi')\langle|C_{\ell m}|^2\rangle.\end{aligned} \tag{14}$$

By changing the order of taking the sum with respect to $\ell$ and $m$ using $n$ having the relation $\ell = n+m$, the expression of eq.(14) turns out to be:

$$\begin{aligned}&\langle \delta r(\varphi,t)\delta r(\varphi',t)\rangle\\ &= \sum_{n=1}^{\infty} 2\big(1-\exp(-\Gamma_n t)\big)\big(N_{n,0}P_n(0)\big)^2\langle|C_{n,0}|^2\rangle.\\ &+ \sum_{m=1}^{\infty}\sum_{n=1}^{\infty} 4\big(N_{m+n,m}P_{m+n}^m(0)\big)^2\big(1-\exp(-\Gamma_{m+n}t)\big)\\ &\quad\times\langle|C_{m+n,m}|^2\rangle\cos(m(\varphi-\varphi')).\end{aligned} \tag{15}$$

where $N_{\ell m}$ is $N_{\ell m} = \sqrt{\frac{2\ell+1}{4\pi}\frac{(\ell-m)!}{(\ell+m)!}}$ and $P_\ell^m$ is the associated Legendre polynomial. By comparing eq.(13) and eq.(15), and using the eq.(7), $\langle|\Delta B_m(t)|^2\rangle$ is found to be expressed as

$$\begin{aligned}\langle|\Delta B_m(t)|^2\rangle = &\sum_{n=1}^{\infty}\frac{2r_0^2k_{\mathrm{B}}T}{\mathcal{E}_{n+m}}(N_{n+m,m}P_{n+m}^m(0))^2\\ &\times\Big(1-\exp(-\Gamma_{n+m}t)\Big)\end{aligned} \tag{16}$$

When nondimensionalizing equation (16),

$$\langle|\Delta\tilde{B}_m(\Delta\tilde{t})|^2\rangle = \sum_{n=1}^{\infty} \frac{2}{\tilde{\mathcal{E}}_{n+m}} \left(N_{n+m,m}\, P_{n+m}^{m}(0)\right)^2$$

$$\times\Big(1-\exp(-\tilde{\mathcal{E}}_{n+m}\Delta\tilde{t})\Big), \tag{17}$$

$$\tilde{\mathcal{E}}_l = \Big(\ell(\ell+1)-2\Big)\Big[\tilde{\kappa}\,\ell(\ell+1)+\tilde{\gamma}\Big] \tag{18}$$

where $\Delta\tilde{B}_m \equiv \Delta B_m/r_0$, $\tilde{\kappa} \equiv \kappa/k_\mathrm{B}T$, $\tilde{\gamma} \equiv \gamma r_0^2/k_\mathrm{B}T$, $\tilde{\mathcal{E}}_{n+m} \equiv \mathcal{E}_{n+m}/\,k_\mathrm{B}\,T$, $\Delta\tilde{t} \equiv \Delta t/t_0$, $t_0 \equiv k_\mathrm{B}\,T/r_0^2\xi = 1/r_0^2\tilde{\xi}$, $\tilde{\xi} \equiv \xi/k_\mathrm{B}T$. Furthermore, when $\Delta\tilde{t} \to \infty$, $\langle|\Delta\tilde{B}_m(\infty)|^2\rangle$ is approximately expressed as

$$\langle|\Delta\tilde{B}_m(\infty)|^2\rangle = \sum_{n=1}^{\infty} \frac{2}{\tilde{\mathcal{E}}_{n+m}} \left(N_{n+m,m}\, P_{n+m}^{m}(0)\right)^2 \tag{19}$$

If we assume that the $n = 1$ mode is dominant, and by introducing two adjust constants $c_1$ and $c_2$, $\langle|\Delta B_m(\infty)|^2\rangle$ in Eq.(16) is approximately expressed as

$$\langle|\Delta\tilde{B}_m(\infty)|^2\rangle \simeq \frac{4}{\pi}\frac{c_1 m + c_2}{m(m+3)\big[\tilde{\kappa}(m+1)(m+2)+\tilde{\gamma}\big]} \tag{20}$$

where $\Delta\tilde{B}_m \equiv \Delta B_m/r_0$, $\tilde{\kappa} \equiv \kappa/k_\mathrm{B}T$, $\tilde{\gamma} \equiv \gamma r_0^2/k_\mathrm{B}T$. $c_1(\tilde{\kappa},\tilde{\gamma})$, and $c_2(\tilde{\kappa},\tilde{\gamma})$. Using fittings, $c_1(1,1) \simeq 1.08$ and $c_2(1,1) \simeq 0$ are estimated. By using eq.(20), the asymptotic behavior of $\langle|\Delta\tilde{B}_m(\infty)|^2\rangle$ when $m \to \infty$ is

$$\langle|\Delta\tilde{B}_m(\infty)|^2\rangle \to \begin{cases} \dfrac{4c_1}{\pi\tilde{\kappa}} m^{-3} & \text{for } \kappa \neq 0 \\[2ex] \dfrac{4c_1}{\pi\tilde{\gamma}} m^{-1} & \text{for } \kappa = 0, \quad \gamma \neq 0 \end{cases} \tag{21}$$

##### J. Fluctuation analysis of condensate surface

To detect the experimental and numerical condensate interface, we consider a polar coordinate system $(r,\varphi)$ from the center of mass of a condensate. The angle $\varphi$ is then discretized by $N$ points separated by a constant angle $\Delta\varphi(= 2\pi/N)$. The camera-pixel-based intensity profile was transformed to the discretized polar coordinate system using the the interp2 function in MATLAB. The condensate contour $r_i(\varphi_i, i = 1, ..., N)$ was measured as the crossing of a global intensity threshold with subpixel resolution using a linear interpolation.

As shown in Fig.1b, the time-averaged shape of the condensate interface is not necessarily a perfect circle. So, we consider the difference in the contour with a time delay $\delta r(\varphi,\Delta t) = r(\varphi, t+\Delta t) - r(\varphi, t)$. Following the Fourier theory, $\delta r(\varphi,\Delta t)$ can be described using the following series;

$$\delta r(n,\Delta t) = \sum_{m=0}^{N-1} \delta r(m,\Delta t) e^{im\varphi_n} \tag{22}$$

here, $\varphi_n = 2\pi n/N$. The fluctuation spectrum can be represented as

$$\delta r(m,\Delta t) = \frac{1}{N}\sum_{n=0}^{N-1} \delta r(n,\Delta t) e^{-im\varphi_n}, \tag{23}$$

which is calculated from experimentally measured $\delta r(n,\Delta t)$ using MATLAB Fast Fourier Transform algorithm. Following Eq. (17), an ensemble average of the fluctuation spectrum $\langle|\Delta B_m(\Delta t)|^2\rangle$ is calculated as

$$\langle|\Delta B_m(\Delta t)|^2\rangle = \frac{1}{M}\sum_{j=1}^{M} |\delta r_j(m,\Delta t)|^2, \tag{24}$$

here, $M$ is the ensemble size. For our analysis, we fit the spectra for modes $5 \leq m < 15$. Modes m = 0 and m = 1 were excluded because they correspond to changes in the average droplet radius and translational motion of the droplet, respectively, and are not relevant to interfacial fluctuation dynamics. Intermediate modes $2 \leq m \leq 4$ were excluded, considering that they may not have fully relaxed within the experimental time window.

#### K. Quantification of concentration ratio

To quantify the concentration ratio of the dilute and dense phases in the nucleus $C_{\mathrm{dil}}/C_{\mathrm{den}}$ after phase separation in the steady state, the condensates were automatically segmented using a single intensity threshold derived from the bimodal fluorescent histogram of the core component or SRRM1-eYFP within the nucleus. To accurately determine concentration, morphological erosion and dilation were applied to the segmented condensates, ensuring that three pixels (approximately 0.48 $\mu$m) near the condensate interface were excluded from the analysis. The concentration ratio, $C_{\mathrm{dil}}/C_{\mathrm{den}}$, was estimated from the average fluorescence intensities of the background-subtracted image in the segmented dilute and dense regions.

#### II. FIGURES

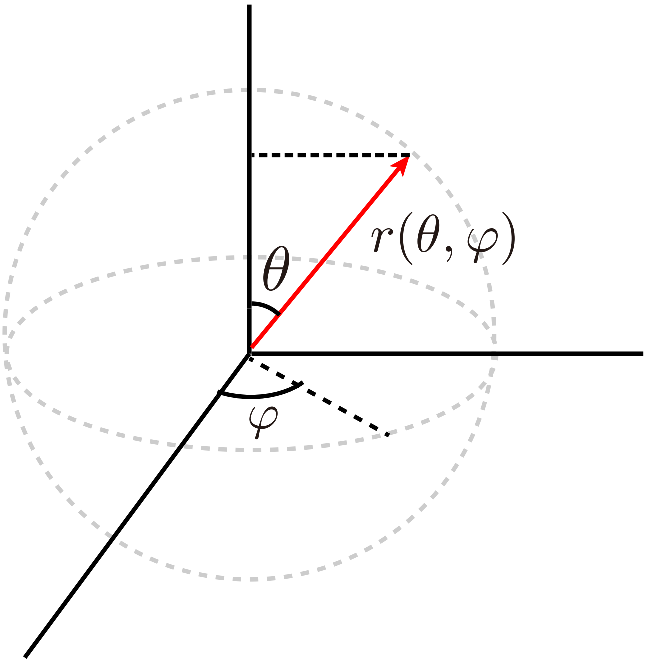


FIG. S1. Representation of a point on the condensate surface using polar coordinates.

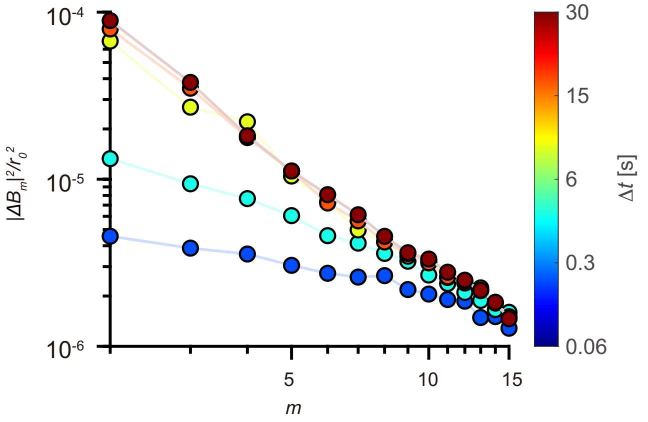


FIG. S2. Time-averaged mode amplitude spectrum $\langle |\Delta B_m|^2 \rangle / r_0^2$ as a function of mode number $m$, evaluated over various time intervals $\Delta t$. Each curve corresponds to a different $\Delta t$. As $\Delta t$ increases, the spectra converge, indicating that $\Delta t = 30$ s provides a good approximation of the infinite-time limit.

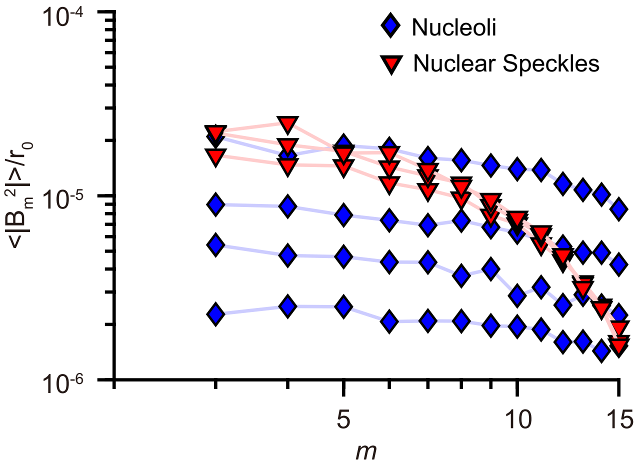


FIG. S3. Noise floor for nucleoli and nuclear speckles. Noise floor with $\Delta t = 750$ sec for nucleoli (4 nucleoli, 2 cells) and $\Delta t = 250$ sec for nuclear speckles (3 nuclear speckles, 2 cells) after fixing the cells with 4 % paraformaldehyde.

[1] D. Bracha, M. T. Walls, L. Avalos, M. tzo Wei, J. E. Toettcher, and C. P. Brangwynne, Mapping local and global liquid phase behavior in living cells using photo-oligomerizable seeds, Cell **175**, 1467 (2018).
[2] S. F. Shimobayashi, P. Ronceray, D. W. Sanders, M. P. Haataja, and C. P. Brangwynne, Nucleation landscape of biomolecular condensates, Nature **599**, 503 (2021).
[3] S. F. Shimobayashi, M. Ichikawa, and T. Taniguchi, Direct observations of transition dynamics from macro-to micro-phase separation in asymmetric lipid bilayers induced by externally added glycolipids, Europhysics Letters **113**, 56005 (2016).
[4] U. Seifert, Configurations of fluid membranes and vesicles, Advances in physics **46**, 13 (1997).
